# Morphological criteria for staging near-hatching embryos of the domesticated Mallard (*Anas platyrhynchos*) and Swan Goose (*Anser cygnoides*)

**DOI:** 10.1101/2024.08.23.609168

**Authors:** Bassel Arnaout, Kaylen Brzezinski, Benjamin Steventon, Daniel J. Field

**Affiliations:** Department of Earth Sciences, University of Cambridge, Cambridge, UK; Department of Genetics, University of Cambridge, Cambridge, UK; Department of Biology, Carleton University, Ottawa, Ontario, Canada; Museum of Zoology, University of Cambridge, Cambridge, UK

**Keywords:** Anseriformes, duck, goose, embryonic development, staging table

## Abstract

Studying waterfowl embryology necessitates reliable and precise staging tables. These are descriptions of embryonic features throughout development from fertilisation to hatching stage, which are used to approximate the extent of embryonic development. Waterfowl staging tables have previously been established based on morphological features from fertilisation to approximately ten days before hatching. Embryonic changes over the final ten days of pre-hatching development have also been documented and proposed as useful staging criteria. However, the reliability of these changes as useful staging criteria across different waterfowl breeds has not been fully examined. To examine the reliability of these criteria— which focus on size changes of the bill and middle toes—for staging near-hatching embryos, we examined twenty-seven Mallard and Swan Geese embryos and measured their beak and toe-length sizes. We compared our results with previously published data, and found that size variation across breeds within the same developmental stage is equivalent to within-breed variation across different stages, suggesting limited reliability of beak and middle toe length sizes for staging waterfowl embryos. Consequently, we devised novel staging criteria for waterfowl based on four easily measurable morphological traits and show that these criteria allow correct stage identification with over 70% accuracy. Our results highlight the importance of quantifying accuracy for developing reliable embryonic staging tables.

## Introduction

Anseriformes (Waterfowl) is one of the most iconic and economically significant groups of birds, factors that have made them the focus of a wide variety of zoological studies, including investigations of embryonic development. Embryological studies of waterfowl have examined the development of many regions of the avian body including the beak (Brugmann et al., 2010), feet (Zou & Niswander, 1996), feathers (Xu et al., 2007), skeleton (Maxwell, 2008), liver (Fancsi, 1982), and ovaries (Ran et al., 2023). Moreover, ecological and conservation studies have studied waterfowl embryos for understanding nest parasitism and embryotoxicity of common pollutants (Brunstrom, 1988; Hoffman, 1978; Hoffman & Albers, 1984; Johnson et al., 1996). Veterinary studies have also examined waterfowl embryos in investigations regarding the viability of artificial insemination (Stunden et al., 1998) and pathology of avian viruses (Bernáth et al., 2006; Fu et al., 2012).

The wide-ranging relevance of waterfowl embryos across different scientific domains places a premium on establishing reliable and precise normal staging tables. A normal staging table is a description of a species’ typical sequence of embryonic features that appear and change, between fertilisation and hatching (Stern, 2018). Staging tables allow investigators to determine how and when embryonic features and body parts develop. Several waterfowl staging tables have been proposed, most of which are partial (i.e. only covering a portion of development between fertilisation and hatching) and focused only on the early stages of development (Chen, 1932; Dupuy et al., 2002, Lukaszewicz et al., 2017). For Mallards (*Anas platyrhynchos*), Chen (1932) described embryonic development up to twelve hours post-fertilisation, and Dupuy et al. (2002) constructed a partial staging table that extends to 72 hours post-fertilisation. For the Swan Goose (*Anser cygnoides*), Lukaszewicz et al. (2017) constructed a partial staging table that extends to sixteen hours post-fertilisation.

Two complete staging tables have been devised for different breeds of Mallards (Koecke, 1958; Li et al., 2019). Koecke (1958) established a staging table using embryos from the Khaki Campbell and Indian Runner breeds, while Li et al. (2019) investigated embryos belonging to the Jinding breed. Li et al. (2019) also constructed a complete staging table for Swan Geese using embryos of the Huo Yan breed. These staging tables applied distinct morphological features for defining embryonic stages 1 to 39; however, they relied exclusively on measurements of beak and middle toe sizes for the final five stages of development (stages 40-45).

The reliance on organ size for staging embryos may reduce the utility of staging tables, as coeval embryos of different breeds may exhibit different beak and toe sizes (Koecke, 1958; Li et al., 2019). Moreover, investigators might use embryos from unknown or hybird breeds. Therefore, establishing broadly applicable staging tables that overcome these limitations is necessary for examining near-hatching embryonic development in a broader range of waterfowl taxa. To overcome these limitations and benefit future investigations of waterfowl development, the present study proposes novel waterfowl staging criteria based on four morphological traits for near-hatching Mallard and Swan Goose embryos.

## Materials & Methods

### Specimen acquisition

Sixteen Mallard (*Anas platyrhynchos*) eggs, two of which belong to Campbell breed, two to the Runner breed, twelve to either Campbell or Runner breeds, and eleven Swan Goose (*Anser cygnoides*) eggs were obtained from Anglia Waterfowl and Poultry, Ipswich, UK. The eggs were incubated in a Brinsea Ovation 56 incubator set at 37.5°C and 40% humidity. Mallard eggs were incubated for 18 to 26 days, and goose eggs for 20-29 days, to obtain embryos older than stage 39 following Li et al. (2019). Embryos were extracted from the eggs following UK Home Office Regulatory guidelines, fixed in Paraformaldehyde (PFA) overnight, then dehydrated, using an ethanol series, on the next day. Ages of the extracted embryos and their replicates are listed in Tables S1-2. Moreover, two deceased *A. platyrhynchos* embryos and fourteen *A. cygnoides* embryos of unknown age were obtained from the same source and were used to assess the precision of our proposed staging criteria (Tables S1-2).

### Size measurements

Beak and middle toe lengths were measured using digital callipers accurate to 0.2 mm and compared with previously published measurements (Koecke, 1958; Li et al., 2019) to assess the reliability of the use of size measurements for staging duck and goose embryos.

### Morphological criteria for staging embryos

To construct a morphological criterion for staging embryos of each species, detailed observations were made using a light stereomicroscope. Specifically, we focused on observations of the development of the nostril, nasofrontal hinge, ankles, and wing feather tracts. We measured the angle of ankle flexure, that is, the angle between the tibiotarsus and tarsometatarsus, by manually pulling together the medial sides of the right and left feet of the embryo. Afterwards, we aligned the arms of a compass parallel to the left tibiotarsus and tarsometatarsus, then measured the angle of the compass with a protractor. Changes in the transparency of the foot webbing, as described by Caldwell & Snart (1974), were observed by manual separation of the toes and looking through the webbing. Afterwards, the development of these traits was divided into phases.

We constructed the staging criteria for each species by tabulating the age of each embryo and the phase of development of each trait. Embryos of similar ages and phases of trait development were grouped together. After grouping the embryos, the mean or modal phase of trait development was calculated for each group, and a stage number was assigned to each group with their corresponding age range and phases of trait development.

### Staging criteria Precision

Precision of our staging criteria was determined by independent staging of several randomly selected embryos by six independent researchers, three for each species. Before each trial, embryos were placed in containers covered in foil, then shuffled in their placement to ensure randomness of the embryo chosen for staging. Subsequently, during the trial, an embryo was picked at random and the phase of development of each trait within the embryo was determined and tabulated. At the end of each trial, the developmental phases of each trait for each embryo were used to estimate its embryonic stage, or range of stages, based on the staging criteria. At the end of all the trials, the estimated stages for each embryo were compiled and compared. The trials for each species were analysed separately.

Deviations among estimated stages for each embryo were quantified by number and magnitude. The magnitude of deviation was assigned a value of 0.5 when two staging estimates partially overlapped. Estimates deviating by one stage were given a value of 1, estimates deviating by two stages were given a value of 2, and so on. Afterwards, we multiplied the total number of deviations with their respective magnitude to obtain a weighted assessment of deviation for each embryo, and this weighted total was divided by the total number of staging trials and converted to a percentage, dubbed the ‘deviation score’. Finally, the compliment of the deviation score, dubbed the ‘similarity score’, was calculated.

### Specimen imaging

Photographs of the nostrils, nasofrontal hinges, and wing feather tracts were taken at the University of Cambridge Museum of Zoology (UMZC). Photographs were taken using a Canon DSLR with a 1.6x lens mounted on a Leica Z16 APO zoom system. Photographs of the embryos and ankles were taken with a 100 mm lens mounted on a vertical stand. The photographs were taken at multiple focal planes and stacked into images using Helicon Focus (7.5.6 Pro).

## Results

### Beak and middle toe size variation

The mean beak, and middle toe, lengths increased with age in our sample with little variance within each age group in both species (Tables 1 & 2). However, comparisons of our measurements to other studies (Koecke, 1958; Li et al., 2019), which utilised the black khaki Campbell and Jinding Mallard breeds and the Huo Yan goose breed, revealed a great deal of intraspecific variation (Tables 1& 2). Indeed, size differences among coeval embryos from different breeds approximates or even exceeds size differences observed among different age groups within a breed (Tables 1& 2).

**Table 1:** Comparison of beak and middle toe lengths for 18-26 day-old embryos from different breeds of the Mallard duck *Anas platyrhynchos*.

| **This study** | **Koecke 1958\*** | **Li et al. 2019†** |
| --- | --- | --- |
| Campbell and Runner breeds | Black Khaki Campbell<br>breed | Jinding breed |

| Day | BL (mm) | TL (mm) | BL (mm) | BL (mm) | TL (mm) |
| --- | --- | --- | --- | --- | --- |
| 18 | 11.8 ±1.1 | 16 ± 4.7 | 13 | 14.7 | 16.9 |
| 19-20 | 11.7 ±0.8 | 18.8 ±2.8 | - | 15.2 | 19.3 |
| 21-22 | 12.5 ±0.45 | 21 ±2.3 | 15 | 15.4 | 19.7 |
| 23-24 | 12.6 ±1.7 | 20.7 ±3.5 | 17 | 16.5 | 20.6 |
| 25-26 | 13.36 ±0.7 | 24.4 ± 2.3 | - | 17.3 | 22.7 |

\* Taken from Koecke, 1958, †Taken from Li et al., 2019. BL = Beak Length and TL = Middle Toe
Length

**Table 2:** Comparison of beak and middle toe lengths of 19-29 day-old embryos of the domesticated Swan Goose *Anser cygnoides*.

| Day | This study<br>Chinese breed |  | Li et al. 2019<sup>†</sup><br>Huo Yan breed |  |
| --- | --- | --- | --- | --- |
|  | BL (mm) | TL (mm) | BL (mm) | TL (mm) |
| 19 | 15±1.4 | 14±1.4 | 13.1 | 14.5 |
| 20 | 16±1.4 | 17.5±0.7 | 14.2 | 16.8 |
| 21 | 17±1.4 | 19.5±0.7 | 14.9 | 18.3 |
| 24 | 18±0.0 | 28±2.8 | 15.5 | 20.4 |
| 26 | 19.5±0.7 | 27±1.4 | 16.3 | 22.7 |
| 29 | 19 | 26 | - | - |

†Taken from Li et al., 2019. BL = Beak Length and TL = Middle Toe Length

### Discrete morphological embryonic changes

Seven easily observable morphological traits were found to change over time in a broadly consistent manner in waterfowl embryos, suggesting that they may be valuable in the establishment of a consistent staging criteria. These traits are listed below and followed by the staging criteria.

### Nostril development in Mallard

The development of the nostril and the visible components of the nasal cavity are described in lateral view and are divided into four phases. Each phase was given a number and denoted with the abbreviation ‘DN’ for duck nostril.

**DN1:** After eighteen to nineteen days of incubation, the nostrils of most embryos appear rostrocaudally elongate with dorsoventrally deep rostral and caudal ends, yielding a peanut-like shape. The cavity lacks any visible nasal components, but has a blank white surface (Fig. 1, Stage 40 DN1).

**Figure 1:**
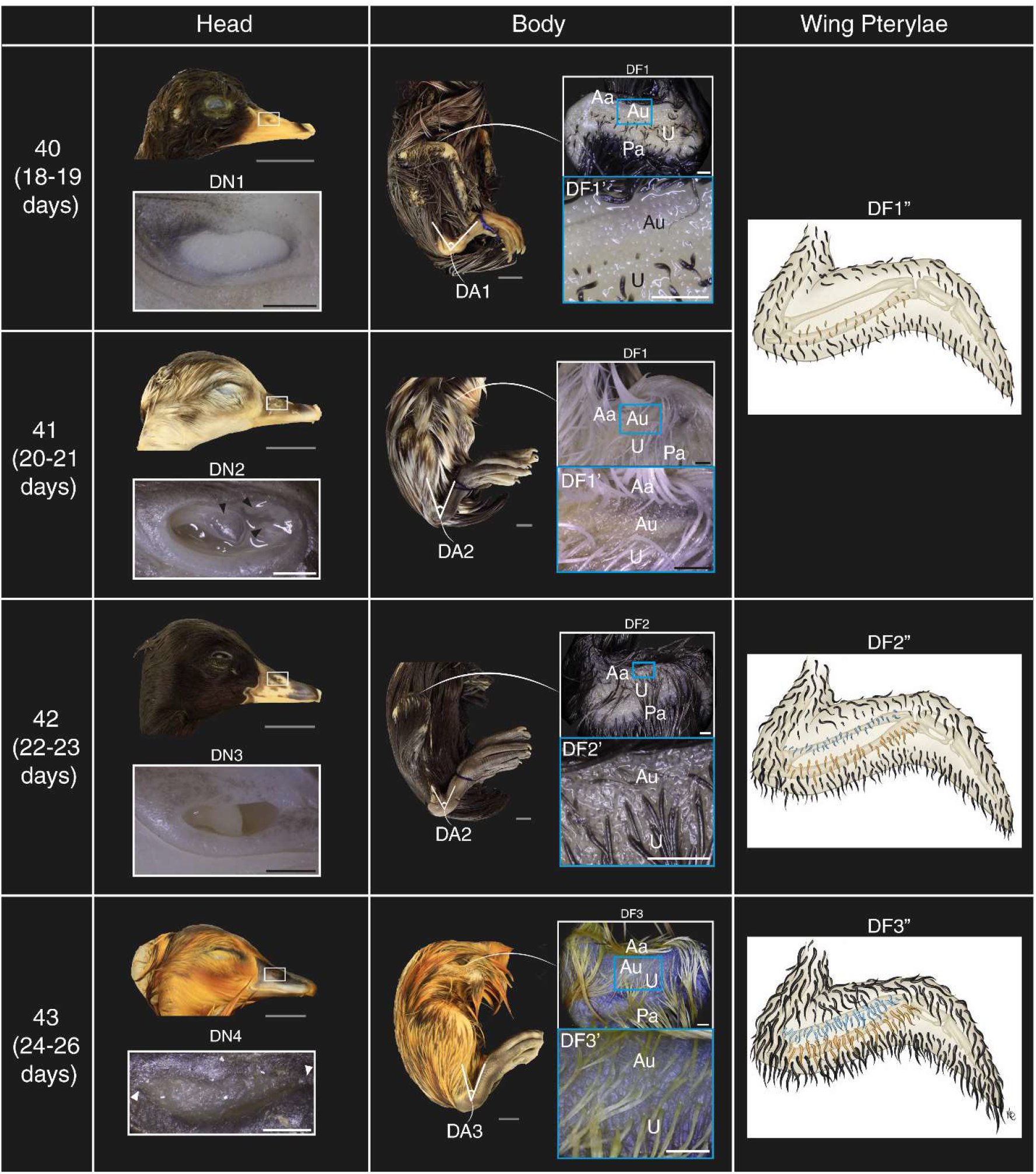
Staging criteria for near-hatching embryos of the Mallard duck (*Anas platyrhynchos*). DN (1-4): Phases of nostril development. DN2: In the second phase of nostril development four nasal components appear, indicated by black arrowheads. DN4: The narrow rostral and caudal ends are indicated by white arrowheads. DA (1-3): phases of changes in angle of ankle flexure. DF (1-3): Phases of feather tract development. DF1’: inset showing the absence of anterior ulnoradial tract at stages 40 and 41. DF2’: inset showing a short anterior ulnoradial tract at stage 42. DF3’: inset showing a longer anterior ulnoradial tract at stage 43. DF (1”-3”): illustrations of the phases of feather tract development. The anterior ulnoradial tract is shown in blue, and the ulnoradial tract is shown in orange. DA (1-3): Phases of ankle development. Grey scale bars are 1 cm and inset scale bars are 1 mm. Abbreviations: Aa; anterior alar, Au; anterior ulnoradial, U; ulnoradial, Pa; posterior alar tracts.

**DN2:** After twenty to twenty-one days of incubation, the overall shape of the nostril appears similar to the previous stage. However, four nostril components appear at this stage: a large triangular caudolaterally-directed component that appears near the caudal end of the cavity, and three triangular rostral components that are directed caudally (Fig. 1, Stage 41 DN2).

**DN3:** After twenty-three to twenty-four days of incubation, the nostril of most embryos is rostrocaudally elongate and is narrower dorsoventrally than in previous stages. Moreover, the three rostral triangular components are no longer visible and only the large caudal component remains visible (Fig. 1, Stage 42 DN3).

**DN4:** After twenty-three to twenty-four days of incubation, the nostril of most embryos is elongated relative to previous stages and takes the form of a slit with pointed rostral and caudal ends (Fig. 1, Stage 43 DN4).

### Nostril development in geese

The development of the nostril and its position within the beak is described in lateral view and is divided into four phases. Each phase was given a number and denoted with the abbreviation ‘GN’ for goose nostril.

**GN1:** After nineteen to twenty days of incubation, the nostril is prominent and is located approximately 1 mm caudal to the back of the egg tooth. The nasal cavity lacks any visible nasal components but is oval in shape and exhibits a blank white surface (Fig. 2, GN1).

**Figure 2:**
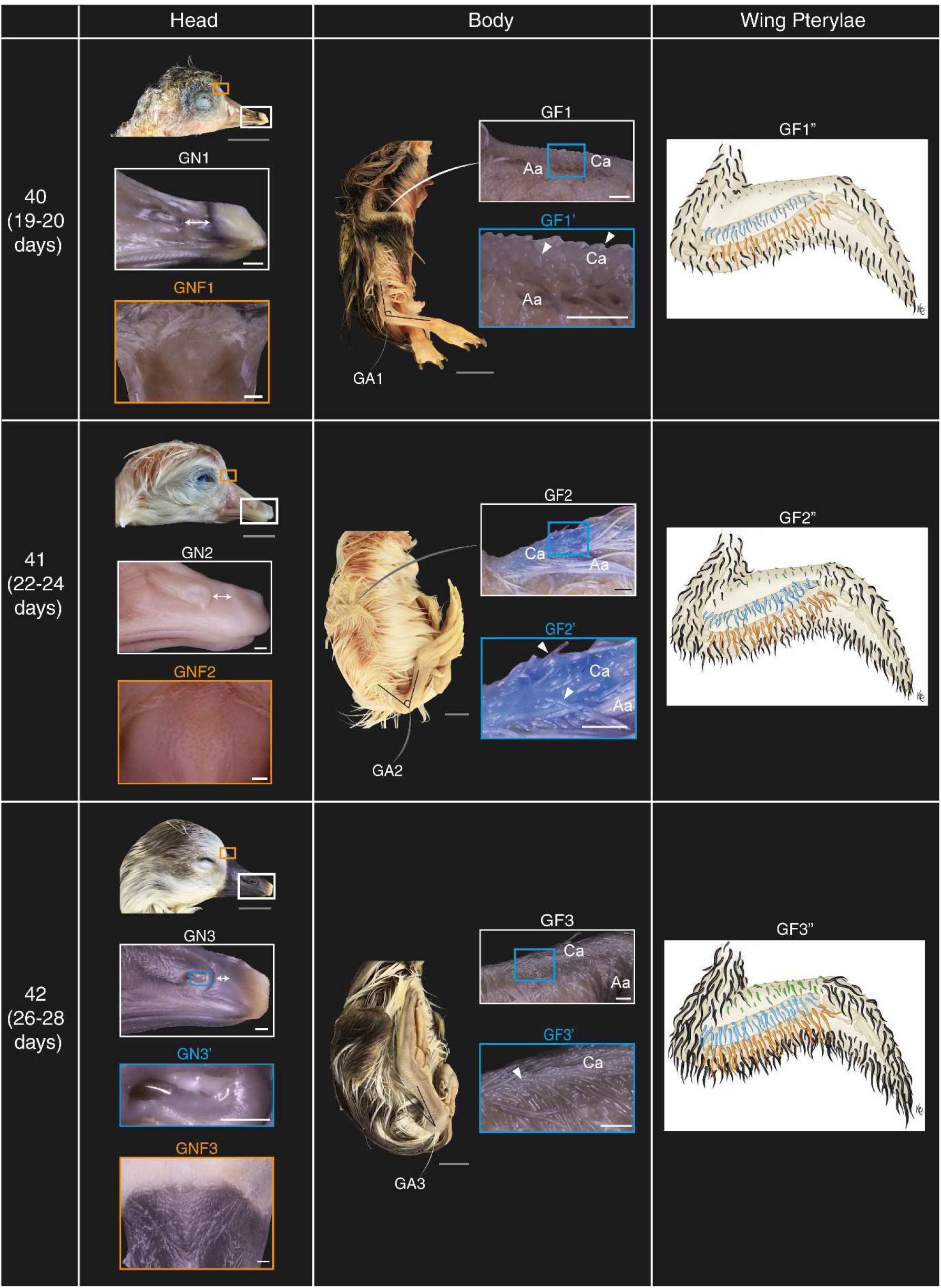
Staging criteria for near-hatching embryos of Swan Goose (*Anser cygnoides*) GN (1-3): Phases of nostril development. In the first and second phases the distance between the nostril and the egg tooth is indicated with a double headed arrow (GN1-2). GN3: At the third phase a pointed white nasal component appears. GNF (1-3): Phases of nasofrontal hinge development. GNF2: One of the deep pits of the second phase of nasofrontal hinge is indicated by a white arrowhead. GF (1-3): Phases of cranial feather tract development; tracts are indicated with black arrowheads. GF (1”-3”): illustrations of the phases of feather tract development. The cranial alar tract is shown in green, anterior ulnoradial tract is shown in blue, and the ulnoradial tract is shown in orange. GA (1-3): Phases of ankle development. Grey scale bars are 1 cm and inset scale bars are 1 mm. Abbreviations: Aa; anterior alar, Ca; cranial alar.

**GN2:** After twenty-two to twenty-four days of incubation, the nostril of most embryos extends rostrally such that it is situated closer to the caudal end of the egg tooth than the previous phase. Moreover, the nostril cavity begins to increase in depth on its rostral side, which gives the nostril a roughly triangular appearance (Fig. 2, GN2). In some embryos, a small light yellow cartilaginous process extends from inside the nasal cavity to the dorsorostral edge of the nostril.

**GN3:** After twenty-four to twenty-nine days of incubation, the nostril of most embryos appears to contact the caudal end of the egg tooth. The nostril cavity acquires a dorsoventrally deeper rostral end and exhibits a more prominent internal cartilaginous process than the previous phase (Fig. 2, GN3 & GN3’).

### Ankle flexure development in mallards and geese

The angle of ankle flexure is measured and described in lateral view. Overall, this angle decreases as development progresses in both species. This process is divided into three phases. Each phase was given a number and denoted with the abbreviation ‘DA’ for duck ankle and ‘GA’ for goose ankle.

**DA1** and **GA1:** After eighteen to nineteen days of incubation, the angle of ankle flexure in most mallards reaches its greatest value (i.e. most obtuse angle), with a mean angle of ∼ 46° (Fig. 1, Stage 40 DA1; Table S1). In geese, after nineteen to twenty days of incubation embryos exhibit a mean angle of 81° (Fig.2, Stage 40 GA1; Table S2).

**DA2** and **GA2:** The angle of ankle flexure in most mallards after twenty to twenty-four days of incubation is smaller than the previous stage, with a mean angle of ∼ 27° (Fig. 1, Stages 41 & 42 DA2; Table S1). In geese, after twenty-two to twenty-four days of incubation embryos exhibit a mean angle of 42.25° (Fig. 2, Stage 41 GA2; Table S2).

**DA3** and **GA3:** The angle of ankle flexure in most mallards after twenty to twenty-four days of incubation is smallest at this stage, with a mean angle of 5° (Fig. 1, Stage 43 DA3; Table S1). As for geese, most embryos after twenty-two to twenty-four days of incubation have a mean angle of 15° (Fig. 2, Stage 42 GA3; Table S2).

### Underwing feather tract development in Mallards

The underwing feather tracts were labelled along the cranial-caudal axis of the wing as the anterior alar, anterior ulnoradial, ulnoradial, and posterior alar tracts, based on Gill (2007). The development of feather tract was divided into three phases, and each phase was given a number and denoted with the abbreviation ‘DF’ for duck feather.

**DF1:** After eighteen to nineteen days of incubation most embryos had developed the anterior alar, ulnoradial and posterior alar tracts. The ulnoradial tract has the smallest feather primordia, and the anterior ulnoradial tract has not yet formed (Fig. 1, Stages 40 & 41 DF1, DF1’, and DF1”).

**DF2:** After twenty to twenty-three days of incubation most embryos exhibit elongated feather primordia of the ulnoradial tract that overlap the posterior alar tract caudally. The anterior ulnoradial tract appears between the anterior alar and ulnoradial tract as small feather primordia that do not extend into the ulnoradial tract (Fig. 1, Stage 42 DF2, DF2’, and DF2”).

**DF3:** After twenty-four to twenty-six days of incubation most embryos exhibit an elongated primordia of anterior ulnoradial tract overlapping the ulnoradial tract (Fig. 1, Stage 43 DF3, DF3’, and DF3”).

### Cranial wing feather tract development in geese

The development of the alar feather tract on the cranial side of the wing was divided into three phases. Each phase was given a number and denoted with the abbreviation ‘GF’ for goose feather.

**GF1:** After nineteen to twenty days of incubation most goose embryos exhibit small feather papilla on the cranial side of the proximal humeral tract (Fig. 2, Stage 40 GF1, GF1’, and GF1”).

**GF2:** After twenty-two to twenty-four days of incubation most goose embryos exhibit very short feather follicles on the cranial side of the proximal humeral tract (Fig. 2, Stage 41 GF2, GF2’, and GF2”).

**GF3:** After twenty-six to twenty-nine days of incubation most goose embryos have longer feather follicles on the cranial side of the proximal humeral tract relative to the previous stage, (Fig. 2, Stage 42 GF3, GF3’, and GF3”).

### Transparency of foot webbing in mallards

The change in the transparency of the webbing between the toes was divided into three phases. Each phase was given a number and denoted with the abbreviation ‘DT’ duck transparency.

**DT1:** The webbing between the toes of most embryos after eighteen to nineteen days of incubation is transparent, such that the shape and colour of objects can be observed through it.

**DT2:** The webbing between the toes of most embryos after twenty to twenty-six days of incubation is less transparent so that only the colours of objects on either side of it are discernible.

**DT3:** The webbing between the toes of most embryos after twenty-four to twenty-six days of incubation is entirely opaque such that neither the colour nor shape of objects on either side of it are discernible.

### Development of the frontonasal hinge in geese

The rostral half of the frontonasal hinge is at the caudodorsal end of the beak and is composed of keratin, lacking any feathers. The development of this region of the hinge was divided into three phases. Each phase was given a number and denoted with the abbreviation ‘GNF’ for goose nasofrontal.

**GNF1:** After nineteen to twenty days of incubation most goose embryos exhibit a smooth, featureless curved surface at the frontonasal hinge (Fig. 2, Stage 40 GNF1).

**GNF2:** The frontonasal hinges of most goose embryos after twenty-two to twenty-four days of incubation have deep pits that lack any specific arrangement (Fig. 2, Stage 41 GNF2).

**GNF3:** Most goose embryos after twenty-six to twenty-nine days of incubation have deep, elongated pits that form a triangular ridge (Fig. 2, Stage 42 GNF3).

### Staging criterion

The phase of development of each trait in each embryo was tabulated, along with the mean and mode phase for each group (Tables S1 & S2). The devised staging criteria consisted of four stages for mallards and three for geese (Figures 1& 2 and Tables 3&4). The precision of the staging criteria was assessed with staging trials (Table S3). The results of the trials revealed a similarity score for the mallard staging criterion of 77% and 74% for the goose staging criterion (Table S4).

**Table 3:**
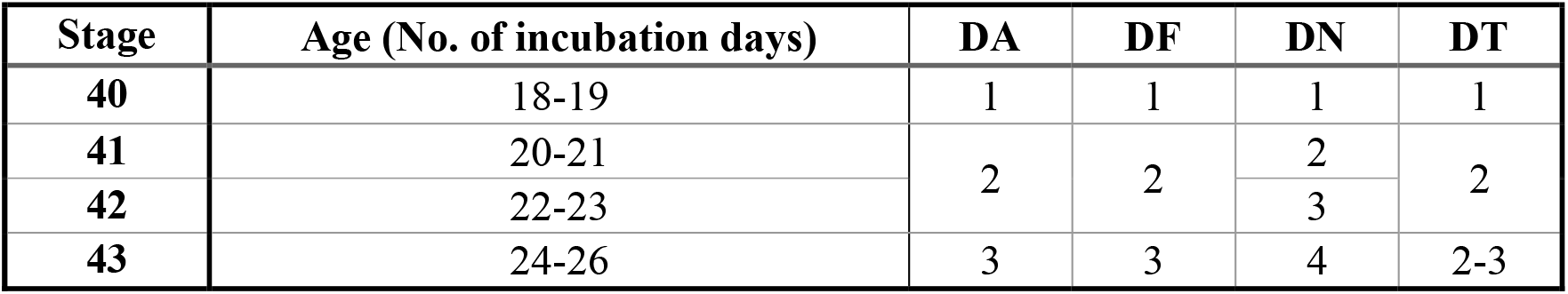
Criterion for staging near-hatching embryos of the Mallard duck (*Anas platyrhynchos*) based on four morphological traits. Abb: A-angle of ankle flexure; F-feather tract; N-Nostril; T-transparency of foot webbing.

**Table 4:** Criterion for staging near-hatching embryos of the domesticated Swan Goose (*Anser cygnoides*) based on four morphological traits. Abb: A-angle of ankle flexure; F-feather tract; N-Nostril; NF-Nasofrontal hinge.

| Stage | Age (No. of incubation days) | GN | GA | GF | GNF |
| --- | --- | --- | --- | --- | --- |
| 40 | 19-20 | 1 | 1 | 1 | 1 |
| 41 | 22-24 | 2 | 2 | 2-3 | 2 |
| 42 | 26-28 | 3 | 3 | 3 | 3 |

## Discussion

Despite the popularity of the use of beak and middle toe sizes in staging near-hatching avian embryos such as chickens (Hamburger & Hamilton, 1992), Japanese Quail (Ainsworth et al., 2010), ducks and geese (Li et al., 2019), these criteria appear to be of limited utility when different or unknown breeds are used. This limitation was apparent from the comparison of the beak, and middle-toe, lengths of duck and goose embryos of the same age range from different breeds (Tables 1& 2). These comparisons revealed that the interbreed variance in beak, or middle toe, lengths of coeval embryos is equivalent to the intrabreed variance in beak, or middle toe, lengths between embryos of different ages (Tables 1 &2). The equivalency between these variances indicated the limited reliability of beak, and middle toe, lengths in staging waterfowl embryos.

The limitations of using size for staging near-hatching embryos illustrate the need for a robust morphology-based staging table. Some of the morphological traits that we used, have been used for staging other bird species; for instance, angle of ankle flexure in embryos of the Society Finch (*Lonchura striata var. domestica*; (Yamasaki & Tonosaki, 1988)). Caldwell and Snart (1974) used the transparency of foot webbing for indexing duck embryos, although they did not establish a staging table. The major difficulty of establishing a high-precision staging table is the considerable degree of intraspecific variation in the rate of development of different morphological traits. To overcome this challenge, we devised a staging criterion that indicates the morphological features that are most likely to be observed for embryos within a particular age range. Consequently, our criteria have three advantages over previously published staging tables for these species. First, our criteria should be applicable to different breeds of ducks, as they were devised using morphological features that commonly occur in two different duck breeds. These morphological features were not discussed in the recent staging table proposed by Li et al. (2019). The use of four morphological features is the second advantage of our staging criterion: Koecke (1958) only mentioned two morphological features rather than the four applied in our criteria. The variability in developmental rate of any individual trait illustrates the benefit of a staging scheme involving multiple traits: the most likely stage for each embryo remains discernable based on the development of all traits considered collectively. The third advantage is the high precision of staging criteria, which ensures the production of consistant staging by different investigators. Consistency in staging embryos is crucial in order to prevent assigning different timings to isochronic developmental features, and events, by different investigators. However, precision of staging tables and criteria is not commonly reported in the published literature

### Interspecific developmental variation

We attempted to construct a comparative staging criteria that would apply for both species, as was carried out by Li and others (2019). Constructing a comparative staging criteria is predicated on the appearance of similar features in different species. Indeed this strategy has been used for comparing early stages of development in waterfowl (Li et al.. 2019) and landfowl (Sellier et al., 2006). However, at near-hatching stages, we could not identify any easily observable morphological traits that are common to both species. The absence of common developmental traits at near-hatching stages indicates an increase in whole-embryo interspecific variability during embryonic development. Furthermore, it is indicative of the necessisty of constructing tissue or organ specific staging tables for comparative developmental analyses.

### Applications of the staging criteria

Our staging criteria can be used for examining duck and goose embryological development at stages close to hatching, which will be beneficial for the exploration of parameters such as osteocranium and chondrocranium development (Maxwell, 2008), feather tract development, foot scale cornification (Koecke 1958), foot webbing pigmentation, wing pterylae, and, tounge (Louryan et al., 2023), liver (Bao et al., 2023), gonads (Mizia et al., 2023), muscle formation (Guo et al., 2021). We hope our approach to staging these taxa will help facilitate a range of investigations into anseriform development in the coming years. We further hope that determing staging precision would become common practice in devising embryonic staging tables or criteria.

## Acknowledgments

We would like to thank Lok Ting Nick New, Delia Capatina, Guillermo Serrano Nájera, Dillan Saunders, Alejandra Guzman Herrera, Oskar Batty, Apolline Delahaye, and other members of the Steventon group for assistance in constructing and verifying the staging criterion and this manuscript. We would also like to thank Natalie Jones (University of Cambridge Museum of Zoology) for facilitating specimen photography. We are thankful to Nick Willis (Anglia Wildfowl farm) for providing duck and goose eggs. This work was funded by UKRI grant MR/X015130/1. For the purpose of open access, the authors have applied a Creative Commons Attribution (CC BY) licence to any Author Accepted Manuscript version arising

## Authors contributions

Bassel Arnaout: Conceptualization; data curation; formal analysis; methodology; writing-original draft; scientific illustrations; writing-review and editing.

Kaylen Brzezinski: providing scientific illustrations

Benjamin Steventon: Conceptualization; formal analysis; funding acquisition; project administration; resources; supervision; writing-review and editing.

Daniel J. Field: Conceptualization; formal analysis; funding acquisition; project administration; resources; supervision; writing-review and editing.

## DATA AVAILABILITY STATEMENT

The data that support the findings of this study are available from the corresponding author upon reasonable request.

