## Supplementary material for "Morphological criteria for staging near-hatching embryos of the domesticated Mallard (*Anas platyrhynchos*) and Swan Goose (*Anser cygnoides*)": Supplmentary table

Supplementary Tables:

| Incubation Day | Replicate Number | Breed | Nostril development | | Angle of Flexure | | Feather track development | | Transparency of foot webbing | |
| --- | --- | --- | --- | --- | --- | --- | --- | --- | --- | --- |
|  |  |  | Score | Mode score | Angle (score) | Mean Angle and mode score | Score | Mode score | Score | Mode score |
| 18 | 1 | NA | 2 | 1 | 19^o^ (2) | 46 ^o^ (1) | 2 | 1 | 3 | 1 |
|  | 2 | Campbell | 1 |  | 35^o^ (1) |  | 1 |  | 1 |  |
|  | 3 | NA | 1 |  | 45^o^ (1) |  | 1 |  | 1 |  |
| 19 | 1 | NA | 1 |  | 57^o^ (1) |  | 1 |  | 1 |  |
|  | 2 | NA | 2 |  | 26^o^ (2) |  | 2 |  | 1 |  |
| 20 | 1 | NA | 1 | 2 | 90^o†^ | 27.5 ^o^ (2) | 1 | 2 | 2 | 2 |
| 21 | 1 | NA | 2 |  | 26^o^ (2) |  | 2 |  | 1 |  |
|  | 2 | NA | 1.5 |  | 34^o^ (2) |  | 2 |  | 2 |  |
| 22 | 1 | Runner | 2.5 | 3 | 20^o^ (2) |  | 2 |  | 2 |  |
| 23 | 1 | NA | 3 |  | 30^o^ (2) |  | 2 |  | 2 |  |
| 24 | 1 | Campbell | 3 | 4 | 34^o^ (2) | 14.7 ^o^ (3) | 3 | 3 | 3 | 2-3 |
| 25 | 1 | NA | 4 |  | 0^o^ (3) |  | 3 |  | 3 |  |
|  | 2 | NA | 4 |  | 8^o^ (3) |  | 3 |  | 2 |  |
|  | 3 | Runner | 4 |  | 9^o^ (3) |  | 3 |  | 2 |  |
| 26 | 1 | NA | 4 |  | 30^o^ (2) |  | 3 |  | 2 |  |
|  | 2 | NA | 4 |  | 7^o^ (3) |  | 3 |  | 3 |  |

**Table S1: Scorings for Mallard duck (*Anas platyrhynchos*) embryos after 18-26 days of development, belonging to the Campbell and Runner breeds, based on four morphological traits.** The variance in scores of the replicates of the same age indicates the large degree of variation in developmental rate in Mallard ducks.

^†^ Excluded for its unusually high value. NA: the exact breed of these embryos is not unknown, but they are most likely to be of the Campbell or Runner breeds.

**Table S2: Scoring for domesticated Swan Goose (*Anser cygnoides*) embryos after 19-29 days of development, belonging to the Chinese breed, based on four morphological traits.**

| Incubation Day | Replicate Number | Nostril development | | Angle of Flexure | | Feather track development | | Nasofrontal hinge development | |
| --- | --- | --- | --- | --- | --- | --- | --- | --- | --- |
|  |  | Score | Mode score | Angle (score) | Mean Angle and mode score | Score | Mode score | Score | Mode score |
| 19 | 1 | 1 | 1 | 90^o^ (1) | 81^o^ (1) | 1 | 1 | 1 | 1 |
|  | 2 | 1 |  | 70^o^ (1) |  | 1 |  | 1 |  |
| 20 | 1 | 1 |  | 84^o^ (1) |  | 1 |  | 1 |  |
|  | 2 | 1 |  | 80^o^ (1) |  | 2 |  | 2 |  |
| 22 | 1 | 2 | 2 | 65^o^ (2) | 42.25^o^ (2) | 2 | 2-3 | 2 | 2 |
|  | 2 | 2 |  | 61^o^ (2) |  | 2 |  | 2 |  |
| 24 | 1 | 2 |  | 25^o^ (2) |  | 3 |  | 2 |  |
|  | 2 | 2 |  | 18^o^ (3) |  | 3 |  | 2 |  |
| 26 | 1 | 3 | 3 | 20^o^ (3) | 15(3) | 3 | 3 | 3 | 3 |
|  | 2 | 3 |  | 16^o^ (3) |  | 3 |  | 3 |  |
| 29 | 1 | 3 |  | 9^o^ (3) |  | 3 |  | 3 |  |

**Table S3: Staging table for Mallard duck (*Anas platyrhynchos*) and Swan Goose (*Anser cygnoides*) embryos at near hatching stages.** Staging was assessed by seven independent researchers on a random selection of embryos. Variable concluded stages are highlighted in bold.

| Species | Sample ID | Age | Trial No. | Trait Scores | | | | Concluded Stage |
| --- | --- | --- | --- | --- | --- | --- | --- | --- |
|  |  |  |  | A | F | N | T |  |
| *Anas platyrhynchos* | D18(1) | 18 | 1 | 2 | 2 | 2 | 2.5 | 41 |
|  |  |  | 2 | 2 | 2 | 2 | 3 | 41 |
|  | D18(2) |  | 1 | 1 | 1 | 1 | 1 | 40 |
|  |  |  | 2 | 1 | 1 | 1 | 1 | 40 |
|  | D21(2) | 21 | 1 | 2 | 3 | 3 | 3 | **42-43** |
|  |  |  | 2 | 2 | 2 | 1.5 | 2 | **41** |
|  |  |  | 3 | 2 | 3 | 3 | 2 | **42** |
|  | D21(1) |  | 1 | 2 | 2 | 2 | 1 | 41 |
|  |  |  | 2 | 2 | 2 | 2 | 1 | 41 |
|  |  |  | 3 | 3 | 3 | 4 or 2 | 3 | **43** |
|  | D22(1) | 22 | 1 | 2 | 2 | 3 | 2 | 42 |
|  |  |  | 2 | 2 | 2 | 2.5 | 2 | 42 |
|  |  |  | 3 | 3 | 3 | 4 | 2 | **43** |
|  | D23(1) | 23 | 1 | 3 | 2 | 3.5 | 2 | **42-43** |
|  |  |  | 2 | 2 | 2 | 3 | 2 | **42** |
|  | D25(1) | 25 | 1 | 3 | 3 | 4 | 2 | 43 |
|  |  |  | 2 | 3 | 3 | 4 | 2 | 43 |
|  |  |  | 3 | 3 | 3 | 4 | 3 | 43 |
|  | D26(1) | 26 | 1 | 3 | 3 | 3.5 | 3 | 43 |
|  |  |  | 2 | 2 | 3 | 4 | 3 | 43 |
|  | AP43?R2 | Unknown | 1 | 2 | 3 | 4 | 3 | 43 |
|  |  |  | 2 | 2 | 3 | 4 | 2 | **42-43** |
|  | AP43?R1 |  | 1 | 2 | 3 | 4 | 2 | 43 |
|  |  |  | 2 | 3 | 3 | 4 | 3 | 43 |
| *Anser cygnoides* |  |  |  | N | A | F | NF |  |
|  | AD40R2 | 19 | 1 | 2? | 1/2 | 1 | 1 | 40 |
|  |  |  | 2 | 1 | 1 | 1 | 1 | 40 |
|  |  |  | 3 | 1 | 1 | 1 | 1 | 40 |
|  |  |  | 4 | 1 | 1 | 1 | 1 | 40 |
|  |  |  | 5 | 1 | 1 | 1 | 1 | 40 |
|  | AD41R2 | 20 | 1 | 2? | 2 | 3 | 2/3 | **41-42** |
|  |  |  | 2 | 1 | 1 | 2/3 | 2 | 40-41 |
|  |  |  | 3 | 1 | 1 | 2 | 2 | 40-41 |
|  |  |  | 4 | 1 | 1 | 2 | 2 | 40-41 |
|  | AD42R2 | 22 | 1 | 1/2 | 2 | 2/3 | 2 | 41 |
|  |  |  | 2 | 2 | 2 | 2 | 2 | 41 |
|  | AD43R1 | 24 | 2 | 2 | 2 | 3 | 2 | 41 |
|  |  |  | 3 | 2 | 2 | 3 | 2 | 41 |
|  | AD43R2 | 24 | 1 | ? | 3 | 3 | 3 | **42** |
|  |  |  | 2 | 2 | 2 | 3 | 2/3 | **41-42** |
|  |  |  | 3 | 2 | 2 | 3 | 2 | **41** |
|  | AD44R1 | 26 | 1 | 3 | 3 | 3 | 3 | 42 |
|  |  |  | 2 | 3 | 3 | 3 | 3 | 42 |
|  |  |  | 3 | 3 | 3 | 3 | 3 | 42 |
|  |  |  | 4 | 3 | 3 | 3 | 3 | 42 |
|  | AD40-41?R3 | Unknown | 1 | 2 | 2 | 1/2 | 2 | **41** |
|  |  |  | 2 | 2 | 1 | 1 | 2 | **40-41** |
|  |  |  | 3 | 2 | 1 | 1 | 1 | **40** |
|  | AD43-44?R2 |  | 1 | 3 | 3 | 3 | 2/3 | **42** |
|  |  |  | 2 | ? | 3 | 2/3 | 2/3 | **41-42** |
|  |  |  | 3 | 2/3 | 2 | 2 | 2 | **41** |
|  |  |  | 4 | 3 | 3 | 2 | 2 | **41-42** |
|  | AD45?R4 |  | 1 | 2/3 | 3 | 3 | 2/3 | 42 |
|  |  |  | 2 | 3 | 3 | 3 | ? | 42 |
|  | AD43-44?R1 |  | 1 | 3 | ? | 3 | 2/3 | 42 |
|  |  |  | 2 | 3 | 3 | 2/3 | 3 | 42 |
|  |  |  | 3 | 3 | 2 | 3 | 3 | 42 |
|  | AD45?R2 |  | 1 | 3 | 3 | 3 | 3 | 42 |
|  |  |  | 2 | 3 | 2 | 3 | 3 | 42 |
|  |  |  | 3 | 3 | 3 | 3 | 3 | 42 |
|  | AD40-41?R1 |  | 1 | 1 | 2 | ? | 2 | **41** |
|  |  |  | 2 | 1 | 1 | 1 | 2 | 40 |
|  |  |  | 3 | 1 | 1 | 1 | 1 | 40 |
|  | AD44-45?R2 |  | 1 | 3 | 3 | 3 | 1 | 42 |
|  |  |  | 2 | 3 | 3 | 3 | 3 | 42 |
|  | AD44?R1 |  | 1 | 3 | 2/3 | 2/3 | 3 | 42 |
|  |  |  | 2 | 3 | 3 | 3 | 3 | 42 |
|  |  |  | 3 | 3 | 3 | 3 | ? | 42 |
|  | AD45?R4 |  | 1 | 3 | 3 | 2/3 | 3 | 42 |
|  |  |  | 2 | 3 | 3 | 3 | ? | 42 |
|  | AD45?R3 |  | 1 | 2/3 | 3 | 3 | 2 | **41-42** |
|  |  |  | 2 | 3 | 3 | 3 | 2 | 42 |
|  | AD43?R4 |  | 1 | 2 | 3 | 3 | 3 | 42 |
|  |  |  | 2 | ? | 3 | 3 | 3 | 42 |
|  | AD44?R2 |  | 1 | 3 | 3 | 3 | ? | 42 |
|  |  |  | 2 | 2 | 2 | 2 | 2/3 | **41** |
|  |  |  | 3 | 3 | 3 | 3 | ? | 42 |
|  | AD40-41?R2 |  | 1 | 2 | 1/2 | 1 | 2 | **40-41** |
|  |  |  | 2 | 2 | 2 | 2 | 2 | 41 |
|  | AD44-45?R1 |  | 1 | 3 | 2 | 3 | 2 | **41-42** |
|  |  |  | 2 | 3 | 3 | 3 | 2 | 42 |

**Table S4: Precision of the proposed staging system for near-hatching embryos of Mallard duck (*Anas platyrhynchos*) and Swan Goose (*Anser cygnoides*).**

| Species | Degree of stage difference | No. of stage differences | Total |
| --- | --- | --- | --- |
| *Anas platyrhynchos* | 0.5 | 3 | 1.5 |
|  | 1 | 2 | 2 |
|  | 2 | 1 | 2 |
|  |  | Weighted total | 5.5 |
|  |  | Deviation Score | 23% |
|  |  | Similarity Score | **77%** |
| *Anser cygnoides* | 0.5 | 14 | 7 |
|  | 1 | 7 | 7 |
|  | 2 | 0 | 0 |
|  |  | Weighted total | 14 |
|  |  | Deviation Score | 26% |
|  |  | Similarity Score | **74%** |
